# MALDI-TOF mass spectrometry and proteomics as phenotypic screening tools for anti-inflammatory drugs

**DOI:** 10.64898/2025.12.01.691706

**Authors:** Leonie Müller, Serena Bateman, Charlie Haslam, Thomas Dawson, Denise Vlachou, Evan Rosa-Roseberry, Lee Booty, Chloe L. Tayler, Andrew Frey, Roland S. Annan, Melanie V. Leveridge, Maria Emilia Dueñas, Rachel E. Peltier-Heap, Matthias Trost

## Abstract

Phenotypic screening is a powerful technology to discover drug candidates in physiologically relevant systems without prior knowledge of molecular targets; however, mass spectrometry (MS) remains underutilised as readout strategy. In this proof-of-concept study, we developed and evaluated two complementary MS-based phenotypic screening approaches to identify anti-inflammatory compounds in human induced pluripotent stem cell-derived macrophages and compared them to a conventional targeted cytokine profiling assay. First, we established a novel MALDI-TOF MS fingerprinting strategy that effectively distinguished macrophage phenotypes, identified phenotype-specific biomarkers, and maintained high-throughput capabilities while reducing cost. Secondly, we performed an in-depth LC-MS proteomic analysis using low cell input on an Evosep-timsTOF HT setup, providing rich molecular detail. Both MS-based approaches demonstrated large comparability with the cytokine assay, with a large proportion of hits overlapping. Notably, the proteomics workflow uniquely enabled deeper insight into inflammation pathway engagement, off-target effects, compound potency, and cytotoxicity. Together, these findings highlight the potential of MS-driven phenotypic screening to enhance early drug discovery by enabling efficient, informative, and cost-effective hit selection.

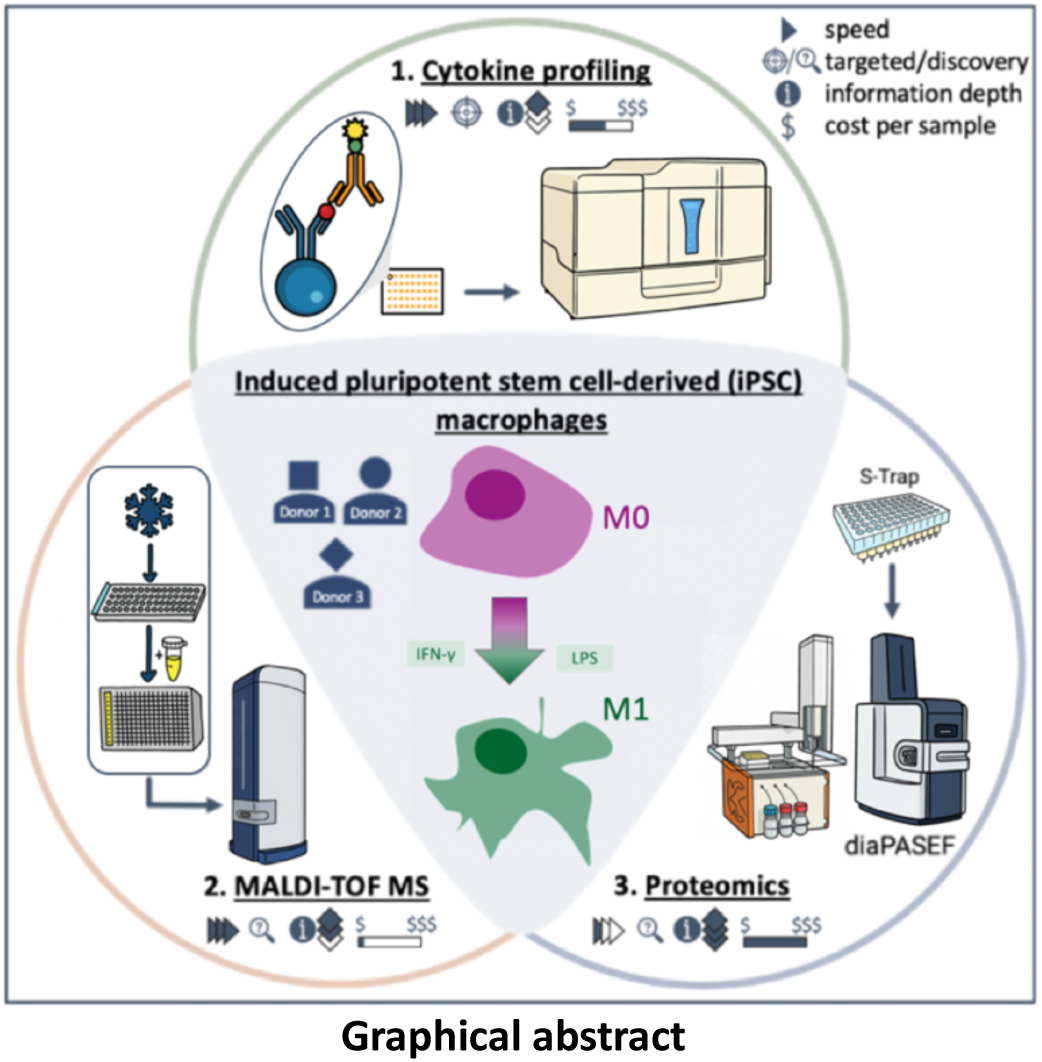

## Introduction

Mass spectrometry (MS) is widely used in drug discovery to analyse disease biology, compound structure/purity, and drug safety (Chen *et al*., 2007; Meissner *et al*., 2022; George *et al*., 2024). However, its application in the lead generation stage remains limited, largely due to historical challenges in throughput and automation. Lead generation typically relies on high-throughput screening (HTS) assays, such as *in vitro* affinity binding and biochemical assays to identify active compounds (hits) from vast libraries (Holdgate *et al*., 2018; Prudent *et al*., 2021). Conventional label-based HTS assays are often limited by high cost, restricted dynamic range, and potential artifacts from non-native substrates. These limitations can be overcome by adopting label-free MS workflows (Leveridge *et al*., 2012; Adam *et al*., 2015). HTS compatible, label-free MS platforms, including RapidFire and MALDI-TOF MS, offer attractive alternatives and have been applied in affinity and biochemical assays (for a detailed review please refer to Dueñas *et al*., 2023).

Yet, most of these target-centric assays fail to capture complex biology, highlighting the value of phenotypic HTS. Phenotypic screens are conducted in physiologically relevant cellular systems, enhancing clinical translatability by revealing both on- and off-target effects (Moffat *et al*., 2017; Vincent *et al*., 2022). However, they require careful design to balance biological complexity and relevance with cost and scalability (Vincent *et al*., 2015). MALDI-TOF MS has emerged as a promising tool for phenotypic screening, offering rapid, unbiased molecular profiling. While widely used in microbial diagnostics (Freiwald and Sauer, 2009), application to mammalian cells is more complex due to their higher molecular and dynamic diversity. Nonetheless, MALDI-TOF MS was successfully used to distinguish cell types (Zhang *et al*., 2006; Karger *et al*., 2010; Petre *et al*., 2020), stress responses (Dong *et al*., 2011; Schwamb *et al*., 2013), differentiation/activation states (Munteanu *et al*., 2012; Blank *et al*., 2022), and to evaluate drug activity (Weigt *et al*., 2018; Weigt *et al*., 2019; Schmidt *et al*., 2024). Broader adoption in drug discovery is hindered by the lack of standardised protocols and data analysis frameworks (Heap *et al*., 2019).

A major challenge in phenotypic screening is the lack of immediate molecular insight, requiring downstream target deconvolution to uncover the target and mechanism of action (MoA) (Moffat *et al*., 2017). While molecular target or MoA knowledge is not mandatory for FDA approval, this information can improve drug efficacy and safety (Becher *et al*., 2016). Advances in proteomics now enable rapid, in-depth profiling, supporting MoA elucidation (Viode *et al*., 2023), off-target identification (Ruprecht *et al*., 2020), and potency assessment (Eckert *et al*., 2024).

Here, we apply MALDI-TOF MS and low-input proteomics to phenotype iPSC-derived macrophages, high-profile drug discovery targets which play a major role in auto-immune and auto-inflammatory diseases, as well as oncology (Wen *et al*., 2021; Bied *et al*., 2023; Zhang *et al*., 2023). Using a dual interferon-γ (IFN-γ) and lipopolysaccharide (LPS) stimulation protocol, a complex multimodal assay was used which modelled robust pro-inflammatory macrophage activation (Bach *et al*., 1996; Pålsson-McDermott and O’Neill, 2004). The performance of MALDI-TOF MS and proteomics was benchmarked against cytokine profiling, a label-based standard HTS method to evaluate macrophage activation (Platchek *et al*., 2020). Building on earlier work using MALDI-TOF MS for immune cell phenotypic drug screening (Marín-Rubio *et al*., 2022), a rapid and cost-effective workflow was developed that distinguished resting from pro-inflammatory induced pluripotent stem cell (iPSC)-derived macrophages across three human donors. Both, MALDI-TOF MS and in-depth proteomics approaches effectively identified phenotype specific biomarkers and enabled identification of anti-inflammatory hits. Proteomics profiling significantly increased the information content for the cell phenotypes, uniquely enabling phenotype-driven compound clustering, as well as insight into inflammation pathway engagement, off-target engagement, and compound potency, highlighting its value for hit selection and MoA studies in lead generation.

## Results

### Established pro-inflammatory cytokine assay identifies 21 inhibitors of inflammation

To test the suitability of novel MS-based screening technologies, we required robust data from a conventional assay platform as a benchmark for inhibitor identification. We deployed an iPSC-derived macrophage model, and a fluorescence-based cytokine profiling assay to quantify four key pro-inflammatory cytokines (TNF-α, IL-6, GM-CSF, and CXCL8). First, differentiated macrophages were treated with a custom compound library of 87 well-annotated chemogenomic library compounds at 10 µM. This library was assembled for hypothesis-driven hit identification, significantly narrowing the scale of the initial screening compared to chemically diverse collections that are utilised in large target-based screens (for compound-target annotation list refer to Appendix table 1). Following three-hour compound incubation, the stimulants (IFN-γ: 20 ng/mL, LPS: 100 ng/mL) were added and incubated for 21 hours before the samples were harvested (Fig. 1A).

**Figure 1.**
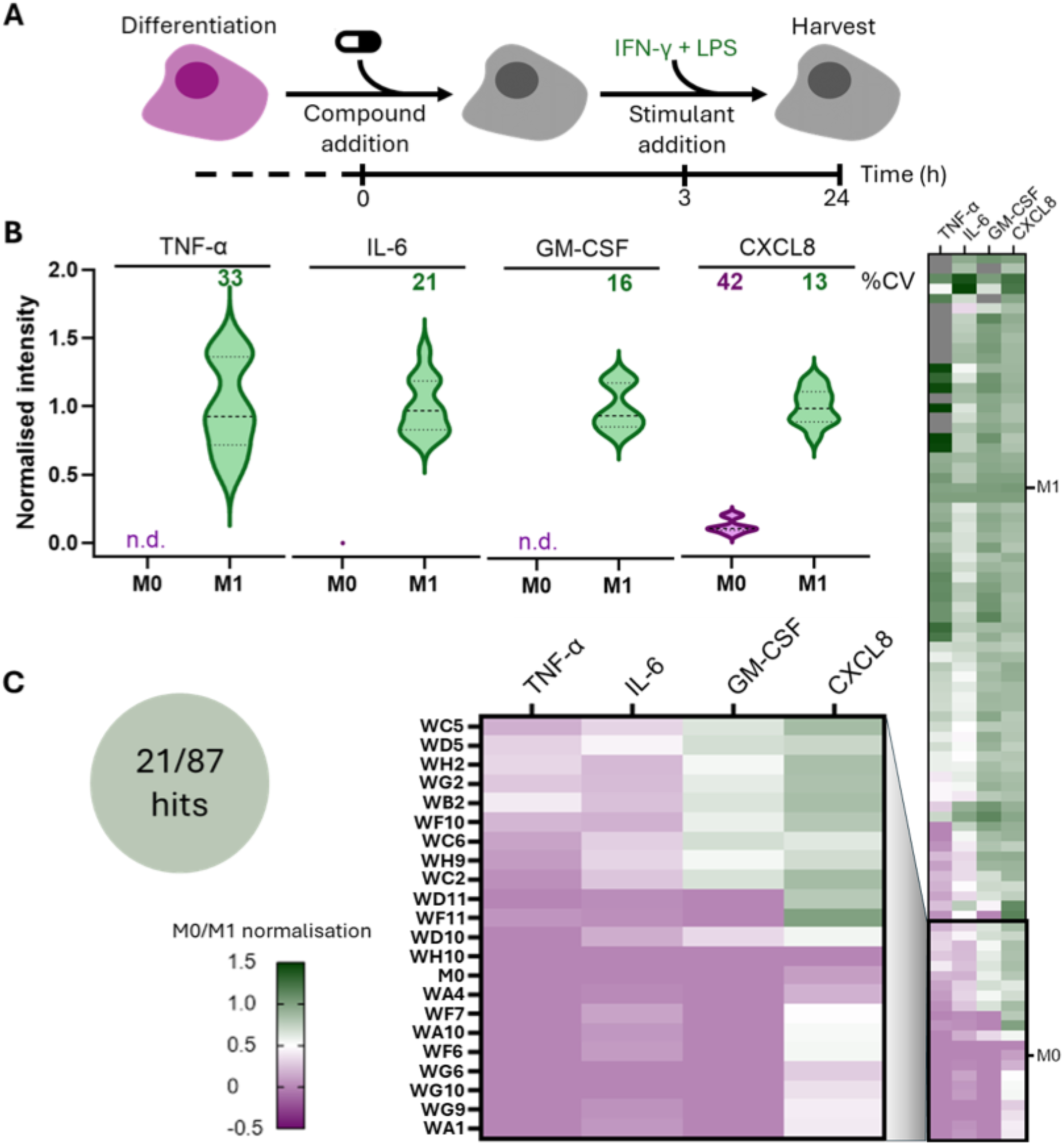
Hit identification in established cytokine profiling compound screen. (**A**) Schematic macrophage treatment and stimulation workflow; six-day macrophage differentiation is followed by compound addition at day seven. After three-hour compound incubation, stimulants (IFN-γ: 20 ng/mL, LPS: 100 ng/mL) were added and incubated for another 21-hours before the samples were harvested. (**B**) Violin plots of the M1 normalised intensities of the cytokines TNF-α, IL-6, GM-CSF and CXCL8 in resting and pro-inflammatory macrophage controls alongside the CV; n_donor_ = 3, n_technical_ = 4 per donor, n_acquisition_ = 1. (**C**) Heatmap of the averaged, M0/M1 normalised cytokine intensities after treatment with 87 compounds. The 21 hit compounds (cut-off: 2-fold standard deviation based on M1 intensity in at least three M1 cytokines) are highlighted in the box; per compound n_cytokine_ = 4, n_donor_ = 3, n_acquisition_ = 1.

The assay demonstrated strong robustness and reproducibility, with coefficients of variation (CVs) ranging from 13% to 33% (Fig. 1B). Of the compounds tested, 21 significantly reduced pro-inflammatory cytokine levels, yielding a 24% hit rate (Fig. 1C). Notably, cytokines such as IL-1β, IL-2, and IL-5 were consistently below the detection limit, highlighting the dynamic range limitations commonly associated with label-based assays (Appendix Fig. 1). In addition to these sensitivity constraints, the high cost of antibodies remains a key drawback of cytokine profiling, despite its widespread use for its speed and reliability, leading us to explore novel MS-based technologies.

### MALDI-TOF MS workflow distinguishes macrophage polarisation states, uncovers polarisation-specific biomarkers, and identifies hits in compound screen

A label-free MALDI-TOF MS-based phenotyping workflow was adapted from published protocols (Heap *et al*., 2019; Marín-Rubio *et al*., 2022), miniaturised, and automated to create a cost-effective assay with throughput comparable to the benchmark cytokine profiling assay (<1 sec per sample) (Fig. 2A). Macrophages derived from iPSCs of three human donors, were treated, stimulated and harvested in 96-well plates following the previously described workflow (Fig. 1A). After mild lysis through freeze-thawing and resuspension, ∼4500 cells per sample were spotted onto a MALDI-TOF MS target plate. After matrix application, metabolite and lipid signatures (*m/z* 400-1,000) were acquired and analysed for mass feature intensities, enabling unsupervised clustering and phenotype biomarker discovery.

**Figure 2.**
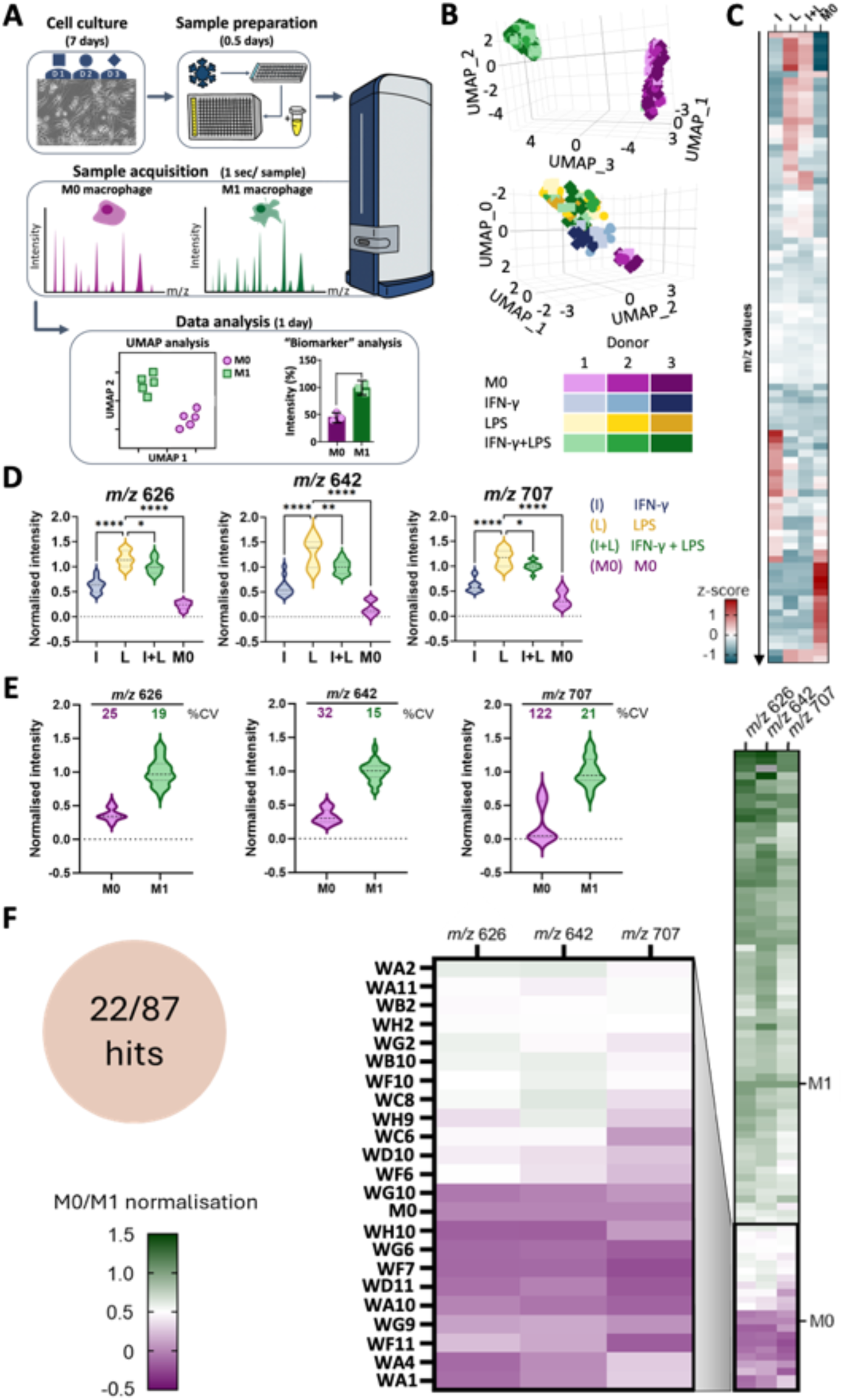
Macrophage phenotype clustering, phenotype-specific biomarker identifications and compound screening by MALDI-TOF MS. (**A**) Schematic assay workflow; iPSC-derived macrophages from three human donors were cultured, treated with compounds, stimulated, and frozen. The cells were thawed and prepared in sample buffer, before being spotted with matrix onto the MALDI-TOF MS target plate. Following acquisition, mass features were extracted, and global phenotype clustering analysed via uniform manifold approximation and projection (UMAP), and phenotype-specific biomarkers identified. (**B**) 3D plot of UMAP analysis showing macrophage polarisation-based clustering across three human donors after dual stimulation with IFN-γ+LPS (n_donor_ = 3, n_technical_ = 46 per donor, n_acquisition_ = 1 per technical) or single stimulation and dual stimulation (n_donor_ = 3, n_technical_ = 4 per donor, n_acquisition_ = 3 per technical), all in comparison to resting macrophages. (**C**) Heatmap displaying the z-scored peak intensities of 95 potential biomarkers which showed differential expression between phenotypes (log2-fold intensity <-0.5 or >0.5). (**D**) Statistical evaluation of intensity differences of selected biomarkers across macrophage phenotypes (One-way ANOVA, Dunnett multiple comparison post-hoc test, * = p-value 0.01 - 0.05, ** = p-value 0.001 - 0.01, *** = p-value 0.0001 - 0.001, **** = p-value <0.0001). (**E**) Violin plots of the M1 normalised peak intensities of the M1 biomarkers *m/z* 626, 642, and 707 in resting and pro-inflammatory macrophage controls within the compound screen alongside the %CV; n_donor_ = 3, n_technical_ = 4 per donor, n_acquisition_ = 3 per technical. (**F**) Heatmap of the averaged, M0/M1 normalised M1 biomarker signal intensities after treatment with 87 compounds. The 22 hit compounds (cut-off: 2-fold M1 biomarker intensity standard deviation in the three *m/z* features) are highlighted in the box; per compound n_m/z_ = 3, n_donor_ = 3, n_technical_ = 1 per donor, n_acquisition_ = 3 per technical.

MALDI-TOF MS data revealed clear phenotype dependent, but donor independent clustering (Fig. 2B). Notably, LPS and IFN-γ+LPS treated macrophages clustered closely together, while IFN-γ only stimulation formed a distinct group, indicating stimulus specific signatures. Visual inspection of z-scored peak intensities for the 95 most discriminatory *m/z* features (log2-fold intensity <-0.5 or >0.5) confirmed the similarity between LPS and IFN-γ+LPS conditions (Fig. 2C), suggesting LPS activation dominates the MS profile. Among these investigated features, several high intensity peaks were identified as potential biomarkers for naïve (M0), IFN-γ, or LPS/IFN-γ+LPS-polarised macrophages (Appendix Fig. 2A). Biomarker intensities were confirmed in the raw spectra (Appendix Fig. 2B). Specifically, *m/z* 626, 642, and 707 emerged as robust pro-inflammatory markers, with maximal intensities under LPS only stimulation, followed by IFN-γ+LPS, and substantially lower signals in IFN-γ and M0 conditions (Fig. 2D).

These pro-inflammatory MALDI-TOF MS biomarkers were subsequently used to identify inflammation inhibitors in the proof-of-principle compound screen. The MALDI-TOF MS readout showed strong reproducibility, with CVs ranging from 15-21% in pro-inflammatory controls, comparable to the cytokine assay benchmark (Fig. 2E). Of 87 tested compounds, 22 significantly reduced the biomarker intensities, yielding a 25% hit rate (Fig. 2F), comparable to the cytokine screen benchmark. These findings demonstrate the potential of MALDI-TOF MS as a rapid, cost-efficient, and robust platform for phenotypic screening and hit triaging.

### Scalar projection enables hit calling from proteomics data

We also wanted to assess the suitability of proteomics for identification of inflammation inhibitors as this technique provided increased biological depth. To ensure sufficient throughput, sample preparation was performed in 96 well plates, and a 60 samples-per-day Evosep workflow (60 SPD = 24 min/sample) with a timsTOF HT instrument operated in diaPASEF mode (injections equivalent to ∼6,000 cells), and data search in DIA-NN against a spectral library (generated from off-line fractioned samples) was used. High quality data was obtained from the 87 compounds in biological triplicates, and 12 controls (4 per donor) per untreated (M0) and IFN-γ+LPS treated (M1) macrophages (288 samples in total). On average 5,153 proteins (±372 proteins) were observed per sample and 5,477 quantified in at least 75% of samples. Workflow reproducibility across donors was indicated by similar mean protein identifications across samples (donor 1 = 5141 ±306 proteins, donor 2 = 5169 ±421 proteins, donor 3 = 5148 ±382 proteins), as well as mean %CV of protein intensities between donors and conditions across the untreated M0/M1 macrophage replicates (10.2 - 13.8%).

Proteomics biomarkers were obtained by comparison between the M1 and M0 macrophage proteome. In total, 45 significantly down- and 217 significantly upregulated proteins were observed (Fig. 3A), including upregulation of M1 cell surface markers (e.g. CD40, CD44, CD48), cytokines (e.g. IL-1α, IL-1β, IL8), IFN-γ stimulated antigen presentation proteins (B2MG, TAP, HLAs), IFN stimulated genes (e.g. ISGs, IRFs, IFITs), and LPS signalling molecules (NFKB1, NFKB2, RELB) (Murugesan *et al*., 2022). To note, LPS pathway activation is mediated via post-translational modifications (PTMs), therefore changes in protein levels are not as prominent as other immune activation pathways (Dunne and O’Neill, 2005; Murugesan *et al*., 2022). Nevertheless, detection of these markers confirms an M1 phenotype, and validates them to probe compound-signalling pathway engagement.

**Figure 3.**
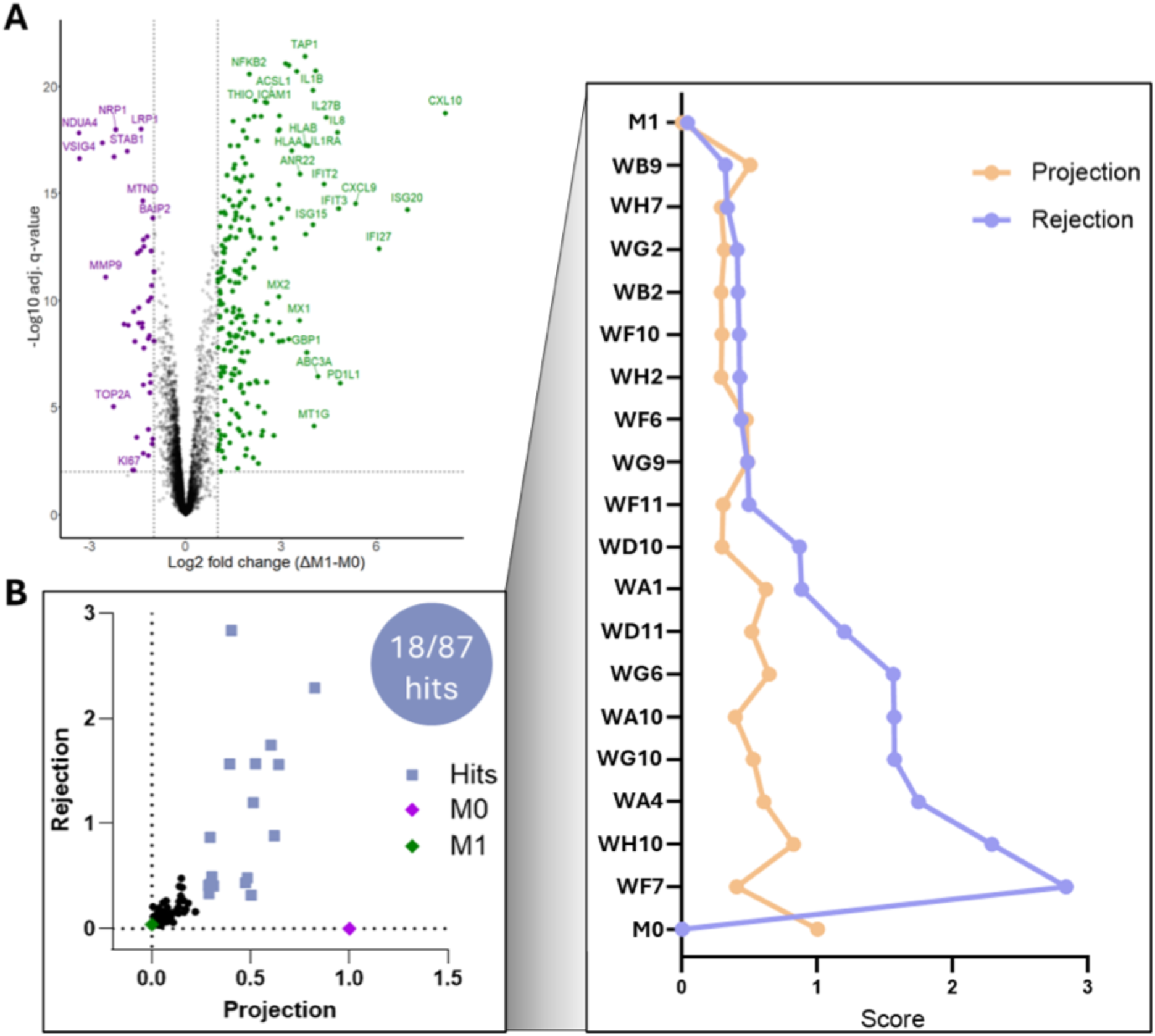
Characterisation of M1 polarisation protein marker and compound screen hits from whole proteome data. (**A**) Volcano plot showing log2 protein fold change (cut-off = 1) against -log10 adjusted q-value (cut-off = 2, i.e. q<0.01) after comparing the M1 macrophages to the resting phenotype with highlighted significantly changed proteins (T-test: Benjamini-Hochberg correction). (**B**) Scatter plot of rejection and projection values obtained by scalar projection analysis of the whole proteome of the controls and 87 compound treatments. Projection and rejection scores of 18 visually distinct hit compounds are highlighted in the box, revealing two subsets based on the magnitude of rejection; per compound n_donor_ = 3, n_technical_ = 1 per donor, n_acquisition_ = 1 per technical.

The proteomics approach utilised complex proteome data instead of individual biomarkers, yet to enable hit calling this high-dimensional data had to be simplified. Here, dimensionality reduction into a 2D model via scalar projection allowed scoring of treatment induced phenotype changes in relation to the controls (Cuccarese *et al*., 2020). In the model, a target axis spans the untreated M1 and M0 samples, allowing scoring for all compounds along this axis (projection), with higher values indicating a shift towards the M0 phenotype. In contrast, high rejection scores (deviation from this axis) often indicate altered biology that is not relevant to the M0 to M1 phenotype axis. Scalar projection distinguished 18 compounds from the main cluster associated with the M1 control (21% hit rate), comparable to the MALDI-TOF MS and cytokine profiling technologies (Fig. 3B). The hit compounds were shifted towards the M0 phenotype, evidenced by projection scores from 0.3 to 0.8 (M0_projection_ = 1), hinting at treatment efficacy. Amongst the hit compounds, two subgroups were distinguished based on their rejection scores (>0.5, n = 9; < 0.5, n = 9), demonstrating that proteomics can indicate compounds that induce off-target effects (high rejection scores) which could affect their therapeutic index.

### Large hit overlap confirms successful hit-triaging via MALDI-TOF MS and proteomics

To evaluate the consistency across screening platforms, we compared compound response profiles obtained from MALDI-TOF MS, proteomics, and cytokine profiling. A moderate correlation was observed between IL-6 cytokine levels and *m/z* 626 MALDI-TOF MS biomarker (r = 0.68, R² = 0.46), the *m/z* 626 MALDI-TOF MS biomarker and proteomics derived projection scores (r = -0.78, R² = 0.61), and lastly proteomics derived projection scores and IL-6 cytokine levels (r = -0.75, R² = 0.57) (Fig. 4A), suggesting broadly concordant phenotypic responses across all three assays.

**Figure 4.**
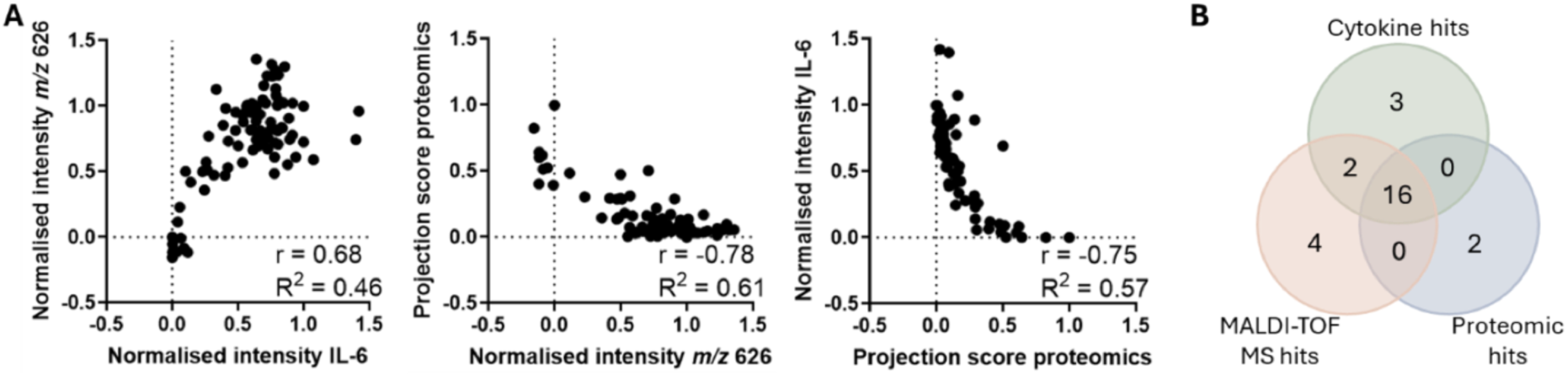
Comparable compound scoring between the three screening technologies. (**A**) Correlation plots of the normalised IL-6 cytokine intensities, normalised *m/z* 626 MALDI-TOF MS biomarker signal intensities, and the proteomics projection scores; correlation analysis. (**B**) Venn diagram of hits identified in the cytokine, MALDI-TOF MS and proteomics screen.

Strikingly, 16 compounds were identified as hits across all platforms (total: cytokine profiling = 21, MALDI-TOF MS = 22, proteomics = 18), highlighting the robustness and reliability of each hit triaging method (Fig. 4B). Two hits (WH9 and WC6) were shared only between the MALDI-TOF MS and cytokine screens, but not detected by proteomics, potentially representing false negatives in the proteomics screen. This could be due to a detection bias in favour of compounds interfering with the LPS pathway as strong LPS signalling biomarker association was observed in MALDI-TOF MS, and significant cytokine secretion is linked to LPS signalling (Pålsson-McDermott and O’Neill, 2004).

Further, several platform-specific hits (cytokine profiling = 3, MALDI-TOF MS = 4, proteomics = 2) were identified. Unique hits from the cytokine and MALDI-TOF MS screens exhibited weak biomarker changes and were likely false positives, a common finding in HTS (Müller *et al*., 2023). These can typically be filtered out through follow-up dose-response assays (Müller *et al*., 2023). In contrast, the two proteomics-specific hits, WH7 and WB9, elicited pronounced phenotypic effects, suggesting they were likely false negatives in the cytokine and MALDI-TOF MS assays. Proteomics appeared particularly sensitive to IFN-γ driven protein level changes, which may explain an enhanced detection of compounds targeting the IFN-γ pathway.

To summarise, the rapid, cost-effective biomarker-based assays prove valuable for initial hit triaging, distinguishing compound activity reliably, and independently from molecular target annotation. Given the concordance between technologies, divergence from the anticipated activity based on the target annotation of each compound may be explained by factors such as compound concentration, or treatment time (Eckert *et al*., 2024). Further, target annotations are derived from diverse *in vitro* and *in vivo* studies, leading to findings which are not always transferrable, emphasising the value of phenotypic screening (Canham *et al*., 2020). However, most phenotypic hit-triaging assays provide limited insight into compound mechanism of action or target engagement. These mechanistic insights can be instead provided by the more comprehensive, albeit lower throughput, proteomics platform.

### Proteomics distinguishes compound-induced inflammation reduction from cytotoxicity

We next wanted to leverage the detailed proteomics dataset to validate inflammation reduction and explore off-target effects. Clustering analysis based on M1 associated protein marker intensities confirmed that only the 18 hit compounds identified through scalar projection analysis exhibited reductions in pro-inflammatory markers (Fig. 5A). Of these, ten compounds induced moderate reductions, while the remaining eight caused pronounced suppression of M1 markers. These eight compounds also displayed high rejection scores (0.9 – 2.8) in the scalar projection analysis, indicating a strong deviation from the M0 and M1 macrophage phenotypes (Fig. 5B). Pathway enrichment analysis confirmed associations with inflammation related pathways, including cytokine signalling in the immune system, interferon signalling, and toll-like receptor cascades across all 18 hits (Fig. 5C). Notably, the subgroup of eight compounds with the highest marker suppression and rejection scores also showed enrichment for the programmed cell death pathway. This indicated that the observed reductions in inflammatory markers were likely driven by compound induced cytotoxicity rather than genuine anti-inflammatory activity.

**Figure 5.**
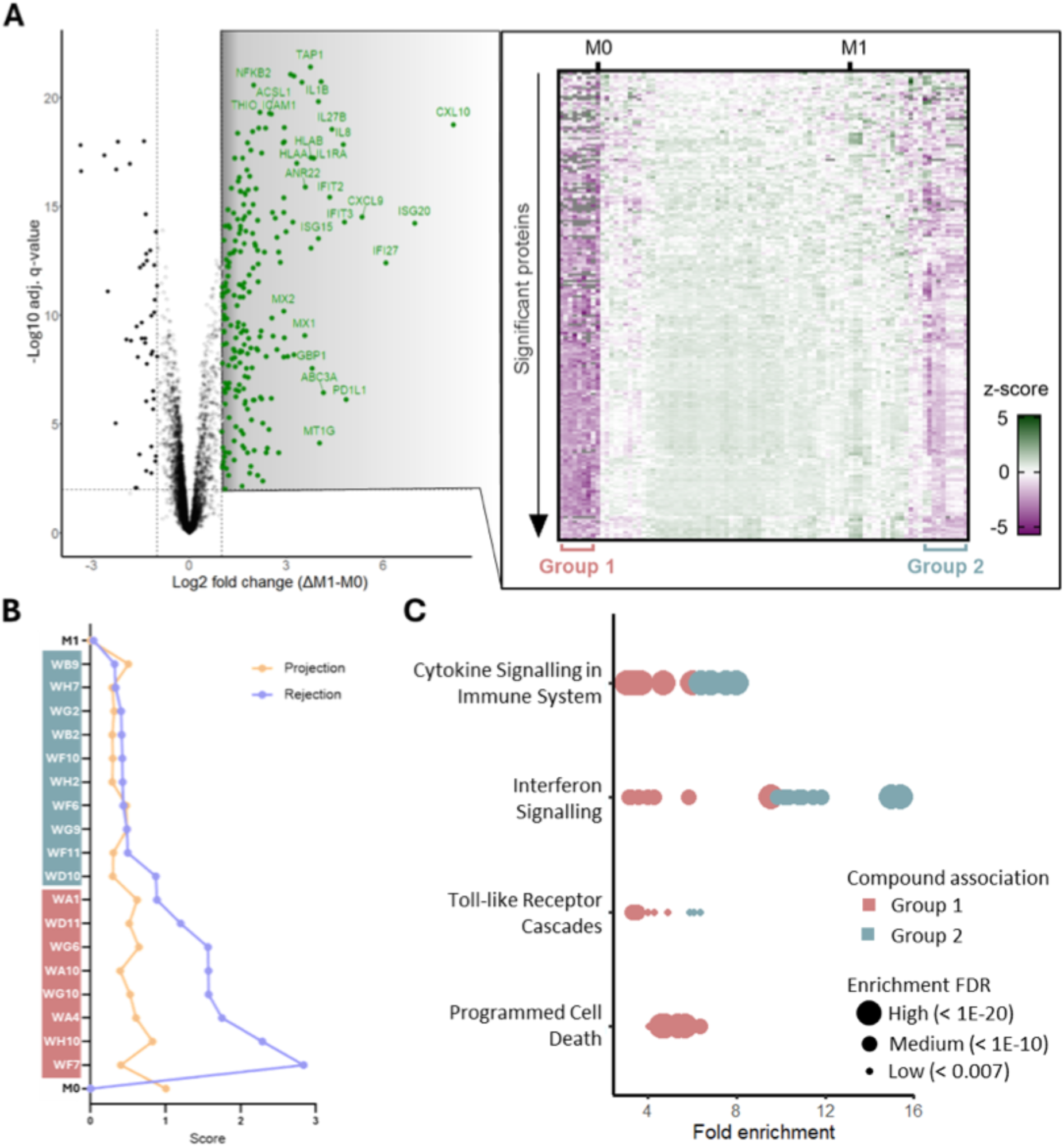
Proteomics confirmed inflammation reduction and uncovered cytotoxicity in a compound subset. (**A**) Hierarchical clustering of z-scored log2-transformed protein intensities of 217 M1 marker proteins (significantly upregulated proteins in the M1 phenotype against the M0 control) across all compound treatments, as well as the controls. Two subgroups were distinguished based on the strength in M1 marker reduction. (**B**) Association of projection and rejection scores identified in earlier scalar projection analysis to the compound subsets identified in A. (**C**) Signalling pathway association analysis (Reactome database) with significantly changing proteins (obtained by comparison against M1 phenotype) in the compound subsets identified in A. Fold enrichment of significantly enriched terms (Enrichment FDR = 0.01 - 1E-20) is plotted.

WA4, WA10, WF7, WG6, WG10 and WH10 treatments were additionally associated with other cellular processes (e.g. protein homeostasis, metabolic reprogramming, cytoskeleton remodelling) and kinases (mean 248 kinases per treatment), indicating that broad cellular processes might be altered due to promiscuous kinase engagement. The compounds are annotated as broad kinase modulators and global kinase modulation is known to reprogram transcription, translation and apoptosis (Sarkar *et al*., 2024). WD11 treatment was also associated with modulation of a large number of kinases, showing kinase promiscuity across its multiple target annotations which include deacetylases, and transcription factors.

In contrast, WA1 activity was not associated with kinases or broader modulated cellular terms. Instead, further inflammation pathway engagement was detected with high significance, suggesting that chronic immune stimulation could be a possible cell death trigger in this case.

These findings underscore the importance of distinguishing true anti-inflammatory effects from off-target cytotoxicity during early drug discovery. Proteomics, through its capacity to capture broad pathway-level changes, offers a critical advantage in advanced hit triaging by identifying compounds that reduce inflammation via undesirable mechanisms such as cell death.

### Proteomics reveals phenotype-driven clustering of non-toxic hit compounds

Next, the phenotypes of the ten non-toxic hit compounds which form three visually distinct compound clusters in principal component analysis (PCA), indicating divergent compound-induced proteomic responses, were further deconvoluted (Fig. 6A). Hierarchical clustering utilising only the 217 previously established M1 protein markers supported this grouping, indicating distinct patterns of inflammation pathway engagement among the groups (Fig. 6B).

**Figure 6.**
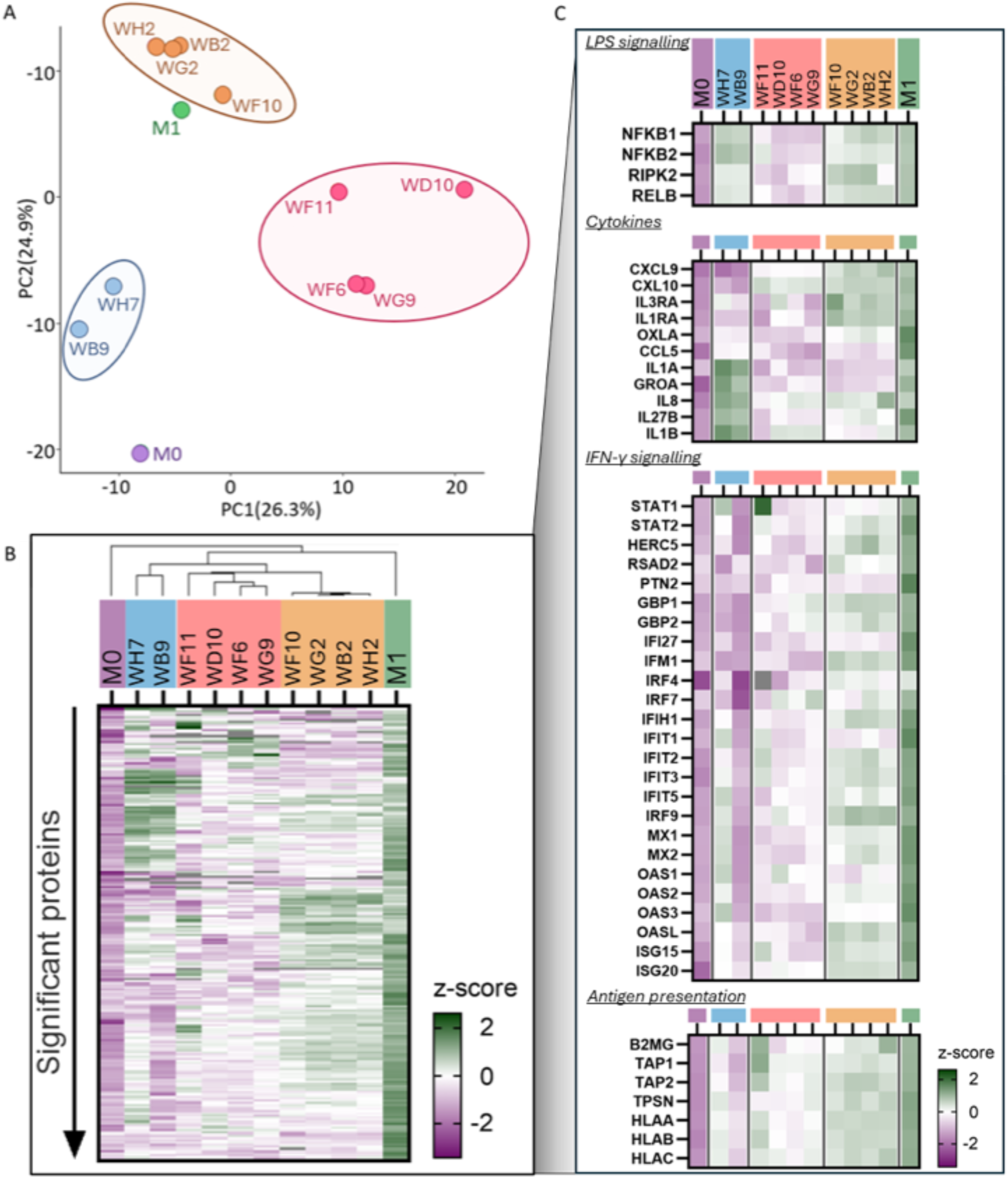
Proteomics guided compound grouping according to inflammation pathway engagement. (**A**) Principal component analysis with whole proteome data from the non-toxic hit compounds and the controls. (**B**) Hierarchical clustering of z-scored log2-transformed protein intensities of 217 M1 marker proteins (significantly upregulated proteins in the M1 phenotype against the M0 control) across the then compound treatments, as well as the controls. (**C**) The M1 marker protein expression analysed in B was investigated in more detail by attributing the proteins to different signalling pathways based on the Reactome database to assess compound engagement in different pathways.

Group 1 consisting of WH7 and WB9, displayed the most prominent shift to an M0 phenotype. Separation from groups two and three of the non-toxic hit compounds was driven by the reduction of several cytokines (e.g. CXCL9, CXCL10, IL1RA) and, more notably, suppression of IFN-γ induced proteins, and antigen presentation (Fig. 6C, Appendix Fig. S3). These are hallmarks of potent JAK inhibitors (De Vries *et al*., 2019), aligning with the compound target annotation. The absence of further kinase engagement suggested pathway specificity, however larger pathway changes upon WB9 treatment indicated differences in drug potency.

Group 2 comprised WF11, WD10, WF6, and WG9. These compounds induced a broad reduction in M1 marker proteins (Fig. 6C), and showed in total 404 significantly altered proteins across treatments (WF11 = 158, WD10 = 248, WF6 = 116, WG9 = 127, Appendix Fig. S4), suggesting interference with fundamental cellular processes. To understand the differences between these compounds, the significantly regulated proteins that were unique within each treatment were analysed. In WF11 treatment, these proteins were associated with nucleus and nuclear lumen localisation, suggesting interference with transcriptional processes. Indeed, the compound is annotated as inhibitor of BRD4, a key transcriptional regulator that impacts a broad set of programs, including inflammation (Lin *et al*., 2022). WD10 treatment associated with kinase activity, suggesting pathway modulation through promiscuous kinase activity. Compound treatment affected fewer kinases than the broad kinase inhibitors which can explain the lack of cytotoxicity. WF6 treatment showed mild association with ubiquitin binding terms in accordance with its deubiquitylase UPS7 target annotation. Further, mild serine-type endopeptidase activity was recorded which explained the tight clustering with the serine/cysteine protease annotated compound WG9, suggesting that both compounds acted via similar molecular pathways. It has been shown that modulation of protease activity such as UPS can influence macrophage polarisation (Xu *et al*., 2020).

Finally, Group 3 consisted of WB2, WF10, WG2, and WH2 which clustered together because a comparable number of significantly altered proteins were observed across the treatments (177 ±9 proteins) out of which only a small portion were treatment unique (7 ±3%; Appendix Fig. S5). The majority of these proteins were upregulated proteins that showed significant association with inflammation terms, explaining only the mildly reduced M1 biomarkers (Fig. 6C), and the clustering closest to the M1 condition (Fig. 6A). Interestingly, all compounds are annotated to mTOR and PI3Ks which are signalling in the PI3K/Akt/mTOR phosphorylation cascade, thereby modulating many cellular processes (Yu and Cui, 2016). Consequently, mTOR interfering compounds are frequently detected hits in phenotypic screens, suggesting unspecific compound activity in this case (Vincent *et al*., 2020).

Together, these results demonstrate that proteomics is a valuable tool to determine cell phenotype-based clustering, uncovering differential pathway engagement upon compound treatment. Moreover, proteomics demonstrated potential to evaluate compound potency and identify potential off-target pathways within the drug discovery pipeline.

## Discussion

This study demonstrates the value of mass spectrometry (MS)-based approaches in early drug discovery, outlining both their advantages and limitations. The MALDI-TOF MS-based metabolite and lipid profiling screen provided a cost-effective, high-throughput alternative to cytokine assays, while capturing rich, phenotype-relevant information. This contributes to the growing number of MALDI-TOF MS applications in eukaryotic cell phenotyping and further validates its utility as a robust tool for hit triaging.

The potential for broader application of this technology is considerable. MALDI-TOF MS has already been used to characterize 66 distinct cell lines and to identify anti-inflammatory compounds active in human monocytes (Karger *et al*., 2010; Marín-Rubio *et al*., 2022). To facilitate adoption across diverse cell types and disease models, a standardised workflow will be essential for reliable technology transfer. Optimising the assay may also involve complementing loss-of-function biomarkers (reflecting reduced pro-inflammatory signals) with gain-of-function markers, to better differentiate true anti-inflammatory effects from cytotoxicity driven reductions. Recent advances in data analysis are further expanding the utility of MALDI-TOF MS. The newly released software *M2ara* enables feature scoring and robustness evaluation, supporting more informed feature selection (Enzlein T *et al*., 2024). Alternatively, clustering approaches based on full spectral fingerprints combined with machine learning algorithms may offer deeper phenotypic resolution and improved classification of desired versus undesired compound responses.

While follow-up assays could help further validate compound activity, these come at the cost of reduced throughput and increased resource demands. To overcome this, we leveraged next-generation, higher-throughput proteomics to both triage and profile HTS hits. Although slower than cytokine and MALDI-TOF MS screens, proteomics provided deeper molecular insights. Ongoing innovations, including LC methods with up to 500 samples/day (Guzman *et al*., 2025), and ultra-fast mass spectrometers such as the Orbitrap Astral (Guzman *et al*., 2024), are rapidly increasing the speed and accessibility of proteomics workflows. In this study, a streamlined bottom-up proteomics workflow capable of acquiring 60 samples per day enabled clustering by phenotype, identification of inflammation pathway inference, cytotoxicity detection, compound potency estimation, and off-target effect discovery. Proteomics is rapidly evolving as a tool for mechanism of action and target identification, with applications that include full proteome fingerprinting, structure-activity relationship analysis, and time- and dose-response characterisation. Specialised methods can provide additional depth by probing post-translational modifications or directly identifying drug-target interactions.

In conclusion, this work advances the integration of high-throughput MS technologies into early-stage drug discovery, highlighting their power to refine hit selection and provide mechanistic insights. Future efforts should focus on extending these workflows to non-immunological cell models, thereby evaluating performance, versatility, and transferability across diverse biological contexts.

## Methods

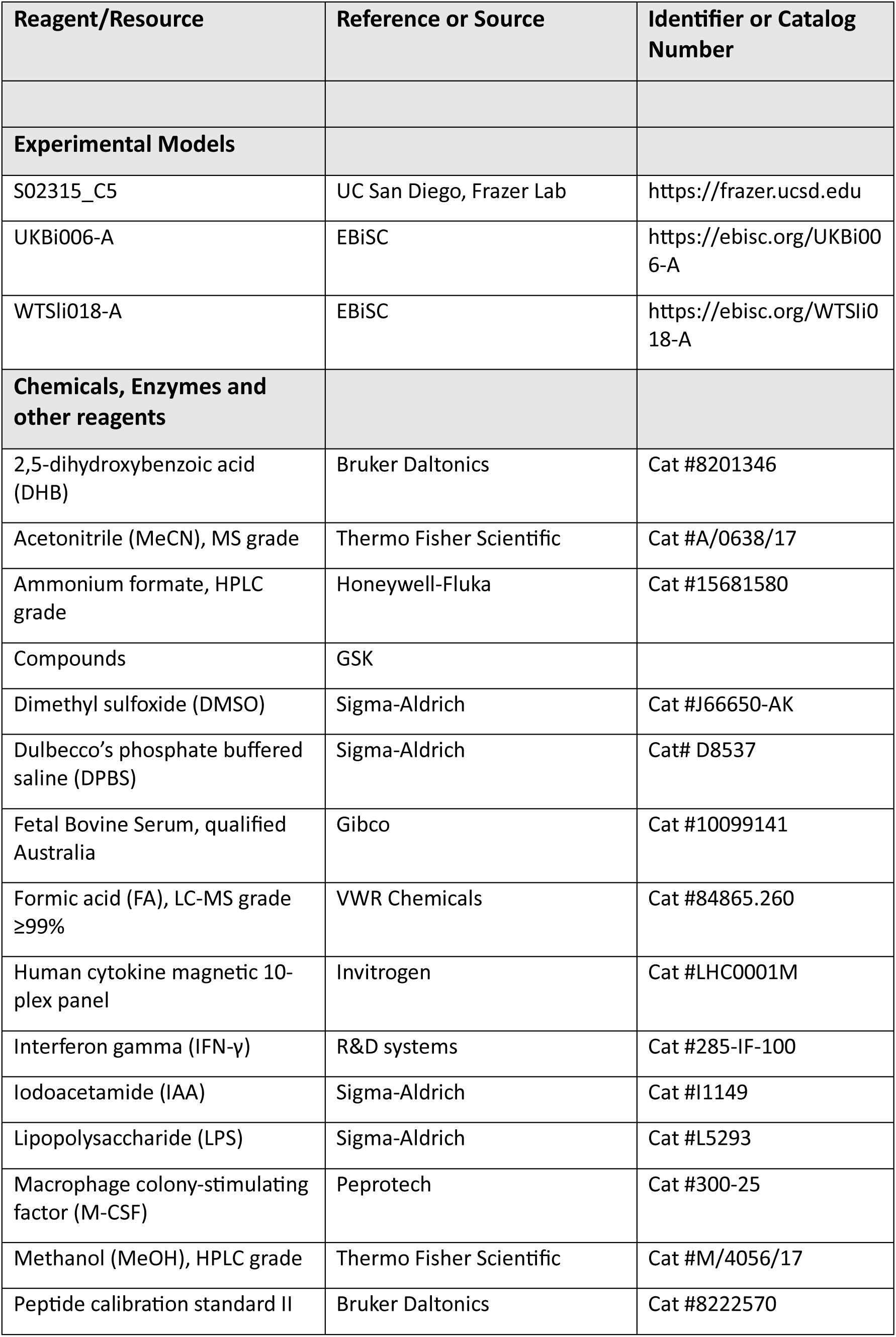
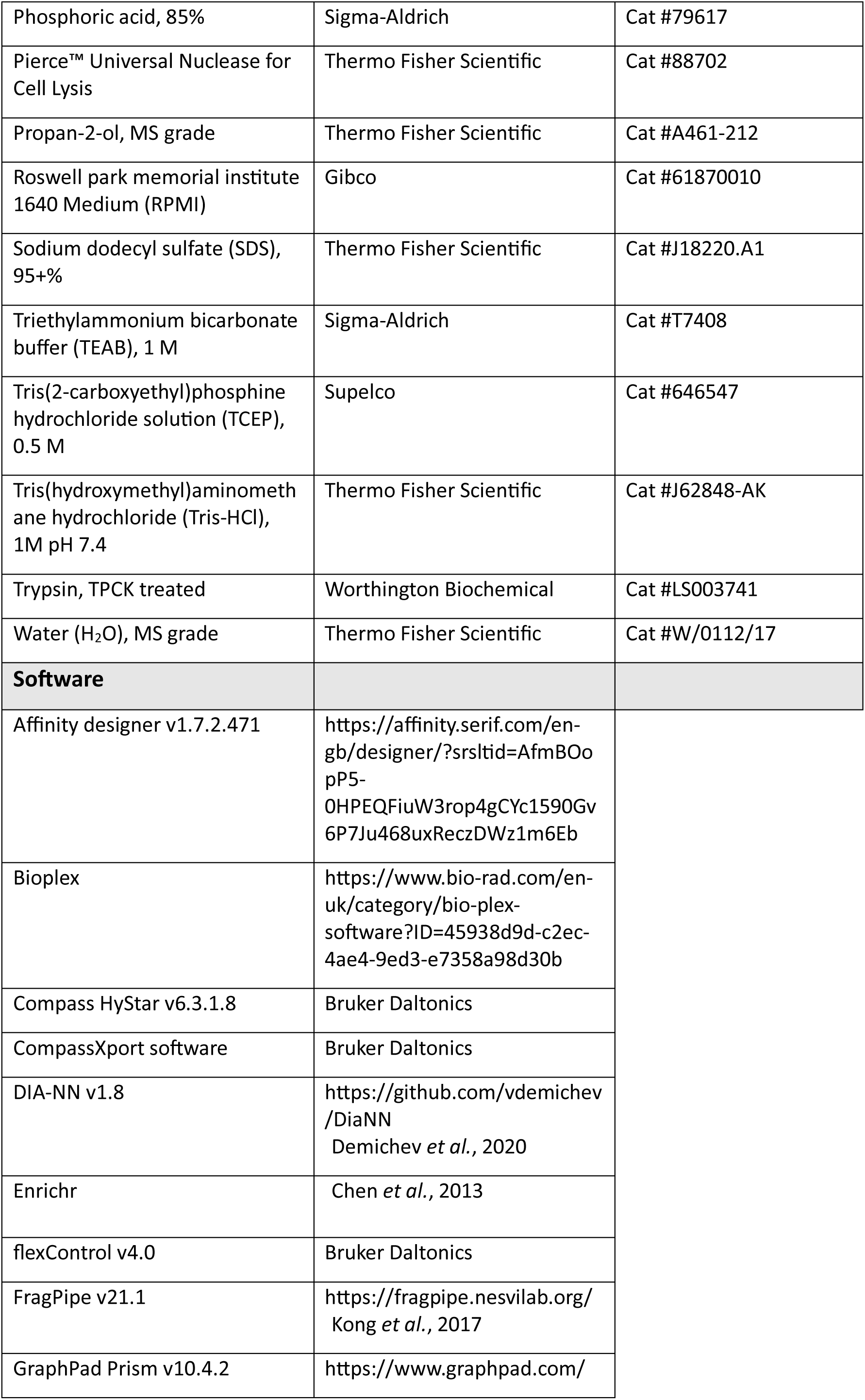
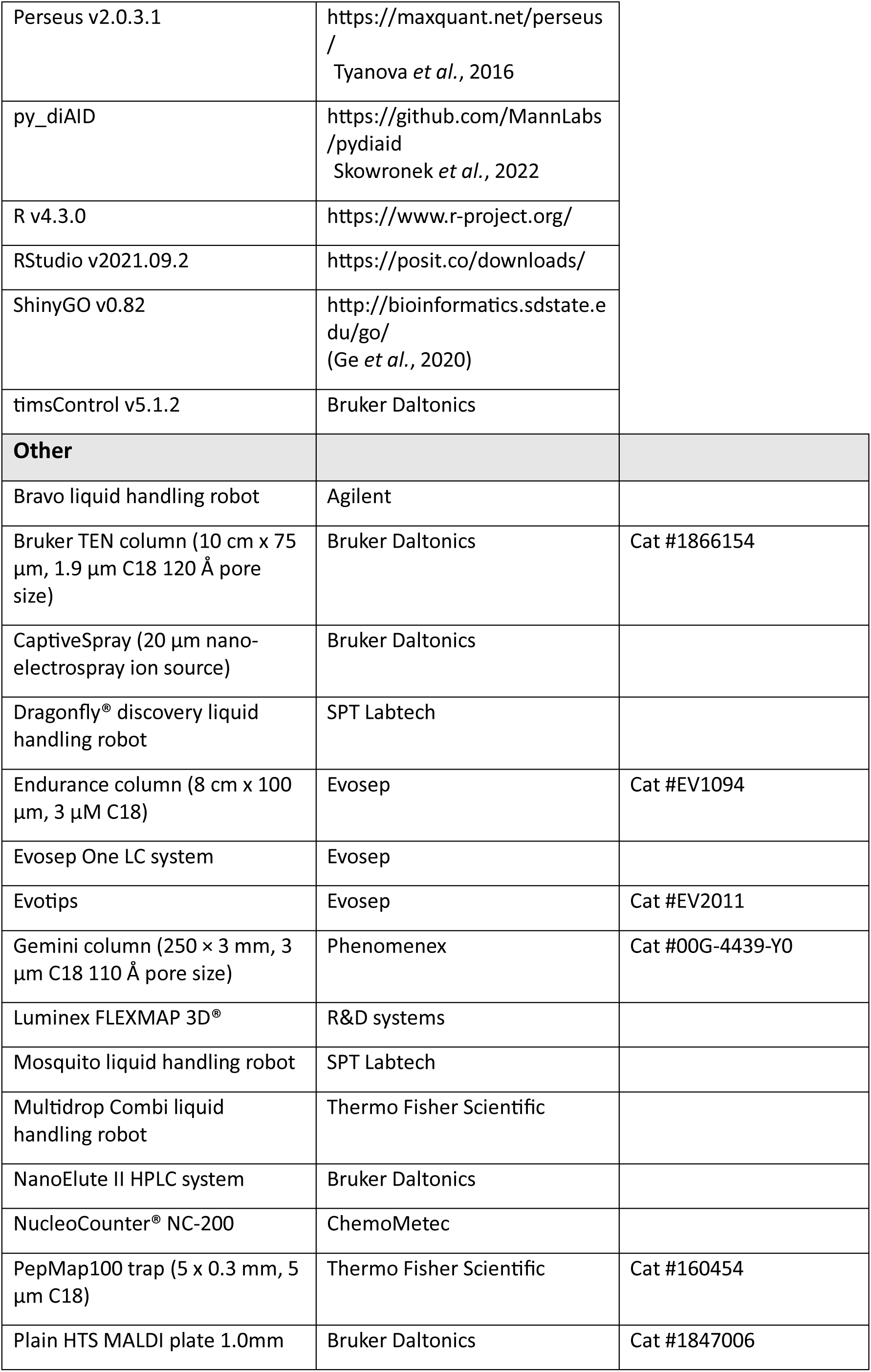
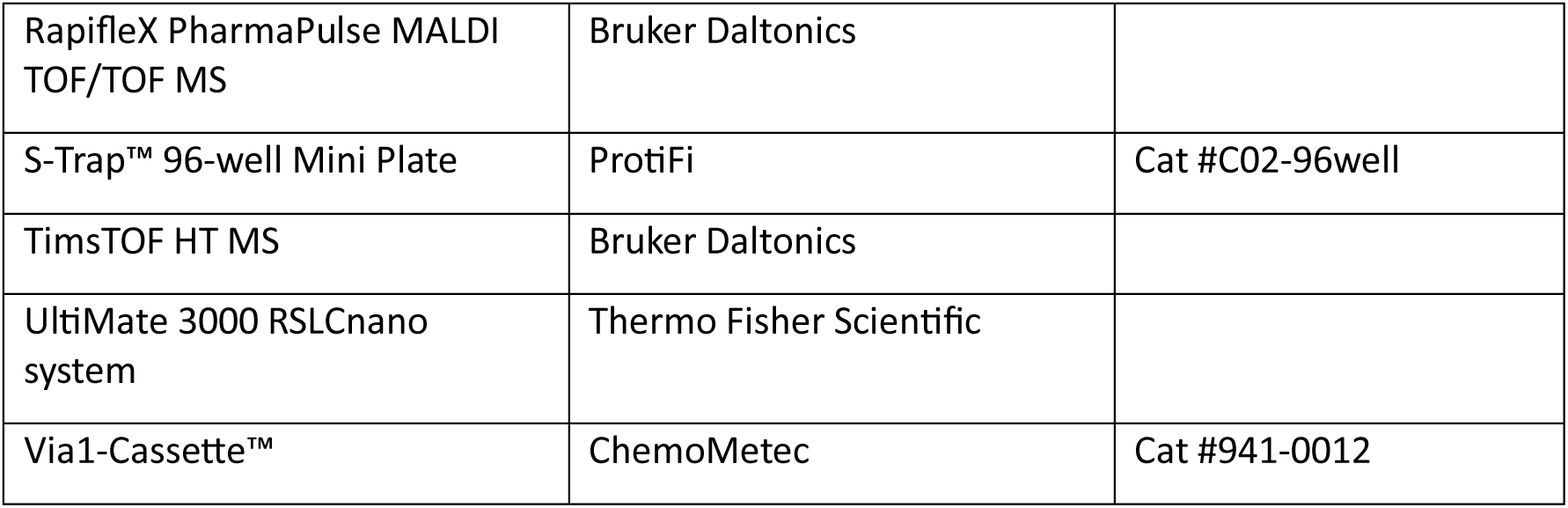

### Cell culture

Detailed protocols outlining the thawing and culture of iPSCs, formation and cultivation of embryoid bodies, as well as monocyte harvest have been previously published (van Wilgenburg *et al*., 2013; Bernard *et al*., 2020). All monocytes were sourced from a GSK proprietary production pipeline, featuring the S02315_C5, UKBi006-A, and WTSli018-A iPSC lines. Monocyte precursors were counted (NucleoCounter® NC-200, Via1-Casette™) and 16.000 cells/well (96 well plate) seeded with a Multidrop Combi reagent dispenser in macrophage differentiation media (RPMI + 10% FBS + 100 ng/mL M-CSF). After 6-day incubation (37°C, 5% CO_2_), the media was replaced and subsequently DMSO (1% final concentration) or compound (10 μM final concentration, Appendix Table 1) added. After a 3 h incubation period (37°C, 5% CO_2_), LPS (100 ng/mL) and IFN-γ (20 ng/mL final concentration) was added to all wells, excluding the unstimulated negative controls. After another 24 h incubation period (37°C, 5% CO_2_), the media was collected, and the cell pellets washed twice with DPBS. All samples were stored at -80°C. Media change, compound addition, stimulation addition, media harvest and cell pellet washes were automated utilising dragonfly® discovery and Bravo liquid handlers.

### Cytokine measurement

The human cytokine magnetic 10-plex panel kit (GM-CSF, IFN-γ, IL-1β, IL-2, IL-4, IL-5, IL-6, IL-8, IL-10, TNF-α) was used. Cell supernatants were thawed over night at 4°C and then diluted in PBS. Bead activation was conducted (50 μL beads/well) and standards prepared in RPMI medium (seven 1:4 dilutions). Standards and samples (50 μL/well) were added to the assay plate and further incubation buffer and assay diluent added (50 μL each). Shaking (30 min, 450 rpm, RT) was followed by incubation over night at 4°C, and again shaking (30 min, 450 rpm, RT). Washing was conducted before the antibody (prepared according to the manufacturer instructions) was added. After shaking (1h, 450 rpm, RT), samples were washed, and Streptavidin-PE added. After shaking (30 min, 450 rpm, RT), and washing, samples were measured by the Luminex FLEXMAP 3D® with the MagPlex™ bead type protocol (7,500 – 15,000 bead region, Bioplex software). The cytokine concentration in the samples was derived from the calibration curves (4/5PL curve fit, Bioplex software) of the standard. A 4/5PL curve fit with extrapolated standard curve data was additionally performed in GraphPad Prism to derive IL-6 and IL-8 concentrations. IL-4 and IL-10 concentrations were not further analysed as they were anti-inflammatory markers and IFN-γ was excluded as it was supplemented to the media for cell stimulation. Next to the standard curve, positive and negative controls were placed on each assay plate to conduct batch correction.

### MALDI-TOF MS sample preparation

The cells were thawed for 20 min at RT. Then, 6 μL/well extraction buffer (100 mM Tris-HCl + 0.1% FA) was added and plates shaken (10 min, RT, 1000 rpm). The cell suspension was transferred into a 384 well plate (#784075, Greiner). After shaking (3 min, 400 rpm), the Mosquito robot was used to conduct three mix cycles (1000 nL volume) in the sample before 1 μL (∼ 4500 cells) was spotted onto a MALDI-TOF MS target plate. The plate was dried before 900 nL matrix (22.22 mg/mL DHB in 70% MeCN + 0.1% FA, 30 min sonicated) was deposited onto the sample with the robot. The plate was dried and then loaded into the RapifleX PharmaPulse MALDI TOF/TOF MS.

### MALDI-TOF MS acquisition

Before sample acquisition, the instrument was calibrated with peptide calibration standard II. Data were acquired in flexControl with an AutoExecute method (positive ion reflector mode, 10 kHz frequency, 10000 shots per spot, M5 custom laser (50 μm x 50 μm scan range), random walk pattern (complete sample), 2000 μm spot diameter, 500 shots per raster position, *m/z* 400 – 1000, mass suppression up to *m/z* 320). The laser power was adjusted to ensure overall spectrum intensity between high 10^5^ to low 10^6^ arbitrary units (a.u.).

### MALDI-TOF MS data analysis

The raw data were converted into .mzxml file format with the CompassXport software to enable peak list extraction in R with the MALDIquant package following a customisable workflow published by Gibb and Strimmer, 2012. The utilised workflow (Appendix Fig. S6) encompassed 1) data import, 2) quality control checks (empty check, irregular check and *m/z* check), 3) square root transformation of intensity, 3) peak smoothing (threshold = 1), 4) baseline correction (TopHat, half window size = 200), 5) intensity normalisation (TIC method), 6) spectra alignment (halfWindowSize = 200, noiseMethod = MAD, signal-to-noise ratio = 7, tolerance = 0.005, warpingMethod = lowess), 7) peak detection, 8) peak binning (strict, tolerance = 0.001), 9) batch correction (limma package, donor based), 10) filtering (excluding peaks with CV >75%, or log2 fold intensity changes between -0.2 and 0.2 based on control M0 and M1 phenotypes), and 11) data visualisation (UMAP generated in R, hierarchical clustering and statistical testing performed in GraphPad Prism). In addition to MALDIquant, the following R packages were required: dplyr, limma, MALDIquantForeign, MALDIrppa, plotly, readMzXmlData, rmarkdown, stats, umap.

### Proteomics sample preparation

Cell culture plates were defrosted at RT for 20 min and subsequently prepared with the S-Trap™ 96-well Mini Plate according to the manufacturer instructions. Briefly, 1x S-Trap lysis buffer (5% SDS + 50 mM TEAB, pH 8.5) was supplemented with Nuclease 5000:1 and 50 μL added per well. Cell lysates were transferred into fresh plates (#EP0030504305, Eppendorf), mixed (15 min, 800 rpm, RT), and centrifuged (1500 xg, RT, 2 min). TCEP (10 mM final concentration) was added and incubated (20 min, 60°C). Then, IAA (10 mM final concentration) was added and incubated (dark, 30 min, RT). Finally, phosphoric acid (1.2% final concentration), and S-Trap binding buffer (100 mM TEAB pH 7.55 in 90% MeOH, 350 μL/well) were added. The samples were loaded onto S-Trap™ plates, centrifuged (2 min, 1500 xg, RT), and washed three times. The digestion buffer (50 mM TEAB pH 8.5) was supplemented with Trypsin (1 μg/sample) and then added and incubated (2 h, 47°C). Peptide elution followed by sequential addition of 50 mM TEAB, 0.1% FA, and 50% MeCN + 0.1% FA with centrifugations (2 min, 1500 xg) between each solvent addition. The samples were dried in a vacuum concentrator.

### Proteomics sample acquisition

Unless otherwise stated, the dried down samples were resuspended in 0.1% FA (20 min, 800 rpm, RT) and acquired with a timsTOF HT MS that was coupled to an Evosep One LC system. The LC system specific C18 Evosep tips were prepared according to manufacturer instructions by 1) 100% MeCN + 0.1% FA wash, 2) isopropanol incubation (5 min), 3) 0.1% FA wash, 4) sample loading (20% sample), 5) 0.1% FA wash, and 6) storage in 0.1% FA. The Evosep system was operated with an Endurance column at 23°C and the 60 SPD (sample per day) protocol (21 min pre-defined gradient, 1 μL/min flow rate). The timsTOF HT was equipped with a CaptiveSpray source which was operated at 1600 V capillary voltage, 3.0 L/min dry gas and 180°C dry temperature. Initially, a pooled sample was subjected to dda-PASEF to generate a custom dia-PASEF method with the py_diAID software. For both methods, general MS (*m/z* 100 – 1700, positive polarity) and TIMS (ion mobility 0.6 - 1.45 1/K_0_, ramp and accumulation times 100 ms) settings were used and in-batch calibration enabled. The dda-PASEF specific MS/MS method settings included: 10 PASEF MS/MS scans, 1.17s cycle time, target intensity 20000, intensity threshold 2500, active exclusion for 0.4 min, *m/z* 150 – 1100, collision energy 20 - 59 eV. The custom 16 variable ion mobility (IM) and mass-to-charge (*m/z)* window pyDIA-PASEF method with two quadrupole positions per window specific MS/MS settings included: *m/z* 300 - 1400, ion mobility 0.6 - 1.4 1/K_0_, cycle time 1.8s, collision energy 20 - 59 eV (refer to Appendix Table 2 for detailed information).

### Extended sample preparation by offine high-pH liquid chromatography fractionation

To generate a custom spectral library for the search of DIA-PASEF acquired samples, offline high-pH liquid chromatography fractionation with subsequent DDA acquisition was conducted. A pooled sample (5% from each sample) was dried in a vacuum concentrator and re-solubilised in 20 mM ammonium formate (pH 8.0). Then, 50% of the pool was separated into 72 fractions with the UltiMate 3000 RSLCnano system that was operated with a Gemini C18 column (72 min linear gradient 1.0% - 37.5% 100% MeCN, flow rate 0.25 ml/min). Fractions were concatenated into 24 fractions (e.g. concatenated 1^st^, 25^th^, and 49^th^ sample), dried in a vacuum concentrator, and resolubilised in 0.1% FA. Acquisition (20% concatenated sample) was conducted with a timsTOF HT that was coupled to a nanoElute II HPLC system which was operated with a Bruker TEN column (60 min linear gradient 0 - 35% 100% MeCN, 300 nL/min flowrate). In addition to the previously described setup, the timsTOF HT MS was equipped a PepMap100 trap and high sensitivity detection was enabled for the DDA acquisition method.

### Proteomics data analysis

Data were searched against the SwissProt *Homo sapiens* database (UP000005640, downloaded in May 2023) with reviewed isoforms (DIA search) or unreviewed isoforms (DDA search), and including Hao laboratory defined protein contaminants (Frankenfield *et al*., 2022). DDA data was searched in FragPipe with MSFragger and Percolator enabled for protein identification. EasyPQP was enabled for spectral library construction, allowing dia-PASEF method construction in py_diAID. The following search parameters were used: mass tolerance for precursor and fragment ions 20 ppm, peptide length 7-50 residues, 1% FDR for peptide- and protein-identification, trypsin specific digestion with 2 missed cleavages, fixed modification (IAA alkylation of cysteine), and variable modifications (oxidation of methionine, acetylation of protein N-termini). DIA data was searched in DIA-NN with the following settings: spectral library, reannotation, trypsin/p specific digestion with 1 missed cleavage, maximum 2 variable modifications, modifications (N-term M excision, C carbamidomethylation, oxidation, N-term acetylation), peptide length 7 - 30 residues, precursor charge 2 - 4, *m/z* 300 - 1400, 1% FDR for peptide- and protein identification, mass and MS1 accuracy 15, MBR, single-pass mode, and QuantUMS. The DIA-NN protein and peptide search output files were processed in R to remove contaminants, exclude proteins with <2 unique peptides, transform (log2), normalise (normalizeMedianAbsValues function, limma package), and batch correct (removeBatchEffect function, donor level, limma package) the data. For statistical testing in R, proteins were filtered (65% valid values in at least one group) and t-tests conducted (limma package, Benjamini-Hochberg). The resulting adjusted p-values and log2 fold changes were displayed as volcano plots in R whilst the list of significantly changing proteins was imported into ShinyGO to perform association analysis with the Reactome database or into Enrichr to perform association analysis with the Kinase Library 2024, and GO Molecular Function databases. Scalar projection analysis was performed as described in Cuccarese *et al*. 2020. Briefly, log2-transformed, normalised, and batch-corrected protein intensities were represented as vectors in an *n*-dimensional space, where *n* corresponds to the number of quantified proteins. A reference (projection) vector was defined as the difference between the barycentre (mean vector) of all M1 and M0 control samples. For each sample, a displacement vector was calculated by subtracting the M1 control mean vector from the sample vector. The projection score was obtained as the scalar projection of the displacement vector onto the reference vector (i.e., the dot product of the displacement and unit projection vector), quantifying movement along the projection axis. The rejection score was defined as the magnitude of the orthogonal component of the displacement vector, corresponding to its residue after subtracting the projection component. Finally, projection and rejection scores were normalised with respect to the control groups (M1_projection_ = 0, M0_projection_ = 1; M1_rejection_ = 0, M0_rejection_ = 0)). A GSK proprietary R script was used to perform scalar projection analysis, and the results visualised with GraphPad Prism. Heatmaps were generated in GraphPad Prism from z-scored and hierarchically clustered data which was obtained from Perseus.

## Data visualisation

FlexAnalysis was utilised for mass spectrum visualisation. Affinity designer, GraphPad Prism, and R were used for workflow and data illustrations. Further, GraphPad Prism was used for correlation analysis.

## Acknowledgements

We thank the members of the GSK screening, profiling, and mechanistic biology (SPMB) team for the initial development of the iPSC-derived macrophage assay and their ongoing support, particularly Tim Ashlin. We would like to thank the GSK iPSC stem cell team for providing the monocyte precursors for this work, particularly Margarida Almeida and Matteo Martufi. We would like to thank Karen Menezes and Amelie Joffrin (GSK, Chemical biology) for providing access to the compounds that were used in this screen. We would like to thank GSK’s AIML Phenomics team for sharing the code implementation used for scalar projection analysis. Furthermore, we would like to thank everyone from GSK and associated companies involved in the sample management supportive of this work.

## Author contribution

L.M.: Designed and conducted most experiments, Data analysis, Manuscript writing, Coding

S.B.: Supported iPSC-derived macrophage culture and automation efforts

C.H.: Generation of iPSC-derived monocyte precursors

T.D.: Generation of iPSC-derived monocyte precursors

D.V.: Supported statistical and scalar projection analysis

E.R-R: Supported scalar projection analysis

L.B.: Conducted cytokine measurements and data analysis

C.L.T.: Conducted cytokine measurements

A.F.: Mass spectrometry maintenance and support

R.S.A.: Funding acquisition, Provided critical feedback

M.V.L.: Funding acquisition, Provided critical feedback

M.E.D.: Conceptualisation, Provided critical feedback, Manuscript writing, Coding

R.E.P.-H.: Conceptualisation, Provided critical feedback, Manuscript writing

M.T.: Conceptualisation, Funding acquisition, Provided critical feedback, Manuscript writing

## Data availability

The mass spectrometry proteomics data have been deposited to the ProteomeXchange Consortium via the PRIDE partner repository (http://www.ebi.ac.uk/pride) with the data set identifier PXDXXXXXX.

